# Natural killer cell activation by respiratory syncytial virus-specific antibodies is decreased in infants with severe respiratory infections and correlates with Fc-glycosylation

**DOI:** 10.1101/750141

**Authors:** Elisabeth A. van Erp, Anke J. Lakerveld, Erik de Graaf, Mads D. Larsen, Rutger M. Schepp, Agnes L. Hipgrave Ederveen, Inge M. L. Ahout, Cornelis A. M. de Haan, Manfred Wuhrer, Willem Luytjes, Gerben Ferwerda, Gestur Vidarsson, Puck B. van Kasteren

## Abstract

**Background:** Respiratory syncytial virus (RSV) is a major cause of severe lower respiratory tract infections in infants and there is no vaccine available. In early life, the most important contributors to protection against infectious diseases are the innate immune system and maternal antibodies. However, the mechanisms by which antibodies can protect against RSV disease are incompletely understood, as both antibody levels and neutralization capacity correlate poorly with protection. We therefore asked whether antibody-mediated natural killer (NK) cell activation correlates with RSV disease.

**Methods:** We performed an observational case-control study including infants hospitalized for RSV infection (n=43, cases), hernia surgery (n=16, controls), or RSV-negative viral respiratory tract infections (n=18, controls). First, we determined RSV antigen-specific antibody levels in infant plasma using a multiplex immunoassay. Subsequently, we measured the capacity of these antibodies to activate NK cells. Finally, we assessed Fc-glycosylation of the RSV-specific antibodies by mass spectrometry.

**Results:** We found that RSV-specific maternal antibodies potently activate NK cells *in vitro*. While the concentrations of RSV-specific antibodies did not differ between cases and controls, antibodies from infants hospitalized for severe lower respiratory tract infections (RSV and/or other) induced significantly less NK cell interferon gamma production than those from uninfected controls. Furthermore, NK cell activation correlated with Fc-fucosylation of RSV-specific antibodies, but their glycosylation status did not significantly differ between cases and controls.

**Conclusions:** Our results suggest that Fc-dependent antibody function and quality, exemplified by NK cell activation and glycosylation, contribute to protection against severe RSV disease and warrant further studies to evaluate the potential of harnessing these activities to develop an effective vaccine.

## Introduction

Respiratory syncytial virus (RSV) is a major cause of severe lower respiratory tract disease in young children, with an estimated 118,000 deaths worldwide in children below 5 years of age annually (1, 2). Despite decades of research, there are currently no market-approved vaccines available against this virus and their development is hampered by the lack of a well-defined correlate of protection. Severe RSV disease is most prevalent in the first six months of life (3, 4), when infants mainly rely on their innate immune system and maternal antibodies for protection against infectious diseases. Considering that maternal vaccination is a potential strategy to prevent RSV disease in infants, it is of pivotal importance to obtain a thorough understanding of the mechanisms by which maternal antibodies might protect against disease, both directly and through their interaction with innate immune cells.

The role of maternal antibodies in RSV infection and disease is unclear. Some studies show that high RSV-specific maternal antibody titers are associated with protection against RSV infection or (severe) disease (5-8). In contrast, other studies do not show a protective effect (9-12) or even indicate an association between high maternal antibody titers and an increased risk of recurrent wheezing (13). Strikingly, the vast majority of studies investigating the role of (maternal) antibodies in RSV infection only look at *in vitro* binding or neutralization capacity, while additional antibody effector functions are not taken into account.

Natural killer (NK) cells are important innate immune cells in the early response to viral infection and their activity is tightly regulated via their interaction with antigen-specific antibodies (14). Engagement of the main NK cell Fc gamma receptor (FcγRIIIa) by antibodies bound to virus-infected cells leads to the release of cytotoxic granules containing perforins and granzymes, a process known as antibody-dependent cell-mediated cytotoxicity (ADCC) (15). In addition, antibody-dependent activation of NK cells is known to result in the secretion of pro-inflammatory cytokines, including interferon gamma (IFN-γ) (16).

Several groups have shown clearance of RSV-infected cells by peripheral blood mononuclear cells (PBMCs) in the presence of monoclonal antibodies, or antibodies from different natural sources, including breast milk, cord blood, nasopharyngeal secretions, and serum (17-21). However, none of these studies shows whether killing was specifically NK cell-mediated. In addition, antibodies from RSV patients and controls have to our knowledge never been compared regarding functional properties other than neutralization, except for antibody-dependent enhancement of infection (22).

In recent years, it has become increasingly clear that the glycan present at the conserved *N*-linked glycosylation site in the Fc-tail of immunoglobulin G (IgG) is variable, and significantly affects antibody functionality (23). Moreover, in some immune responses, such as with HIV (24), dengue (25), and alloimmune diseases (26, 27), reduced fucosylation of the elicited antigen-specific antibodies can affect disease outcome by increasing the affinity for FcγRIIIa. In autoimmune diseases, IgG antibodies can display atypical galactosylation which affects disease severity through mechanisms that are not yet understood (28). The potential impact of IgG glycosylation on (protection from) RSV disease has not been studied to date.

Here, we used samples from an observational case-control study of 84 infants (<7 months of age) that were hospitalized for RSV (cases), an inguinal hernia surgery (uninfected controls), or viral respiratory infections other than RSV (RSV-negative infected controls). We first measured plasma levels of antigen-specific IgG against the RSV attachment (G) and fusion (F) proteins. We then assessed the capacity of RSV-specific antibodies to activate NK cells *in vitro* as measured by CD107a surface expression, which is a marker for degranulation, and IFN-γ production. Finally, we assessed the Fc-glycosylation status of total and RSV-specific antibodies by mass spectrometry. Our findings suggest that antibody-mediated NK cell activation contributes to protection against severe RSV disease and should be further investigated as a potential means to improve vaccine efficacy.

## Methods

### Study design

Plasma samples from hospitalized infants were collected during an observational case-control study in 2010-2014 and have been described before (22). For the current study, hospitalized infants below 7 months of age with PCR-confirmed RSV infections were included as RSV cases. Age-matched infants admitted for inguinal hernia repair surgery were included as uninfected controls, whereas infants admitted to the hospital for viral respiratory tract infections other than RSV were included as RSV-negative infected controls. The ePlex System (Genmark Dx, Carlsbad, CA, USA) was used for identification of respiratory viral pathogens and RSV subtype in nasopharyngeal aspirates. No RSV was detected in uninfected controls and RSV-negative infected controls. Blood samples were collected in heparin tubes within 24 h after admission. Patients with congenital heart or lung disease, immunodeficiency, or glucocorticoid use were excluded. The study protocols were approved by the Regional Committee on Research Involving Human Subjects Arnhem-Nijmegen (serving as the IRB) and were conducted in accordance with the principles of the Declaration of Helsinki. Written informed consent was obtained from the parents of all infants.

### Cells and viruses

Peripheral blood mononuclear cells (PBMCs) were obtained from healthy adult volunteers at the National Institute for Public Health and the Environment (RIVM, the Netherlands). Blood was collected in heparin tubes and the mononuclear fraction was isolated by density gradient centrifugation (Lymphoprep, Nycomed). Isolated cells were cultured in Roswell Park Memorial Institute medium (RPMI, Gibco) supplemented with 10% heat-inactivated fetal calf serum (hiFCS, Hyclone, GE Healthcare), 1% penicillin/streptomycin/glutamine (PSG, Gibco), and 5 ng/mL recombinant human IL-15 (Biolegend). Before use, PBMCs were rested overnight at a density of 1×10^6^ PBMCs/mL at 37°C and 5% CO_2_. Vero cells (ATCC CCL-81) were cultured in Dulbecco’s Modified Eagle Medium (DMEM, Gibco), supplemented with 5% hiFCS and 1% PSG.

Recombinant RSV-X (GenBank FJ948820), used as a coat in the NK cell activation assay, was propagated in Vero cells and purified between layers of 10% and 50% sucrose by ultracentrifugation. The 50% tissue culture infectious dose (TCID50) per mL was determined on Vero cells using the Spearman and Karber method (29). RSV-A2 (ATCC VR1540), used as a coat for the enrichment of RSV-specific antibodies for mass spectrometry, was propagated in Vero cells, purified and concentrated by filtration, and inactivated by the addition of 2% Triton X-100.

### Control sera/plasma

The International Standard for Antiserum to Respiratory Syncytial Virus (16/284, National Institute for Biological Standards and Control) was used as control in the NK cell activation assay and glycosylation study. An in-house adult reference serum pool (RIVM) was used as standard in the multiplex immunoassay. Additionally, cord blood plasma pooled from >10 individual donors was (mock-) depleted for RSV-specific antibodies by incubation with mock- or RSV-infected Vero cells, respectively. Depletion of RSV-specific antibodies was confirmed by multiplex immunoassay as described below. Cord blood samples were collected from umbilical cords of healthy neonates born by cesarean delivery at Radboudumc Nijmegen (the Netherlands). All mothers provided written informed consent.

### RSV-specific multiplex immunoassay

To quantify the concentration of RSV-specific IgG and IgA, a multiplex immunoassay was performed as described before (30). Briefly, diluted plasma samples were incubated with RSV antigen-coupled beads, including G_A_, G_B_, post-F, and pre-F (DS-CAV1 mutant) (31). All antigens were produced in eukaryotic cells. Captured antibodies were detected with secondary R-phycoerythrin-labeled goat-anti-human IgA F(ab’)_2_ (Southern Biotech) or IgG F(ab’)_2_ (Jackson Immunoresearch Laboratories). IgG was measured on a Bio-Plex 200 (Luminex Corporation) in combination with Bio-Plex Manager software version 6.1 (Bio-Rad). IgA was measured on a Flexmap 3D (Luminex Corporation) in combination with Xponent version 4.2 (Luminex Corporation). For each analyte, median fluorescence intensity (MFI) was converted to arbitrary units/ml (AU/ml) by interpolation from a 5-parameter logistic standard curve from an in-house reference serum pool. Whereas all samples were tested in the NK cell activation assay, for some samples not enough volume remained to be tested in the IgG (n=8) and/or IgA (n=11) multiplex immunoassay. These samples were dispersed equally across the 3 different groups.

### NK cell activation assay

Sterile Immulon ELISA plates (ThermoScientific) were coated with intact RSV-X particles at a concentration of 1.7×10^5^ TCID50/well. After overnight coating at 4°C, wells were washed with PBS and blocked with 10% hiFCS in PBS for 30 min at RT. After washing with PBS, coated plates were incubated for 2 hours at 37°C with 10-fold diluted plasma, to allow for antigen-specific immune complex formation. Subsequently, unbound antibodies and other plasma constituents were washed away and 1×10^6^ PBMCs were added per well, together with Brefeldin A (BD Bioscience) and anti-human CD107a-PerCP/Cy5.5 (clone H4A3; BioLegend). PBMCs were incubated with the opsonized virions for 4 hours at 37°C, after which the cells were stained for flow cytometric analysis as described below. Incubation of PBMCs in RSV-coated plates in the absence of antibodies was used to determine the background signal. The PBMC supernatants were stored at −80°C and used for detection of perforin secretion by ELISA (Mabtech). The ELISA was performed following the manufacturer’s protocol and optical density was measured at 450 nm using an ELISA reader (BioTek).

Due to the limited amount of fresh PBMCs per donor, we were unable to test all samples using cells of all 6 donors. For this reason, we could test 63 of the 84 plasma samples on cells from all 6 PBMC donors. For 18 samples, we were able to test with PBMCs from 2-5 different donors. Three samples were only tested on a single donor; these were all RSV-negative infected controls. Excluding these 3 samples had no effect on the subsequent statistical analysis.

### Flow cytometric analysis

PBMCs that were incubated with opsonized virions as described above were stained for flow cytometric analysis. Extracellular staining was performed with anti-human CD3-FITC (clone UCHT1, Biolegend), anti-human CD56-PE (clone HCD56, Biolegend), and Fixable Viability Staining-eFluor780 (eBioscience). After fixation and permeabilization, cells were intracellularly stained with anti-human IFN-γ-PECy7 (clone B27, Biolegend). Flow cytometric analysis was performed using the FACS LSR Fortessa X20 (BD Bioscience). In all experiments, NK cells were gated as the CD3^−^, CD56^+^ population. The gating strategy is depicted in **Fig S1**. FlowJo software V10 (FlowJo, LLC) was used for data analysis.

### Purification of plasma antibodies for glycosylation analysis

Total IgG antibodies were captured from 2 µL of plasma using Protein G Sepharose 4 Fast Flow beads (GE Healthcare, Uppsala, Sweden) in a 96-well filter plate (Millipore Multiscreen, Amsterdam, The Netherlands) as described previously (11). To isolate RSV-specific antibodies from plasma, maxisorp ELISA plates were coated with inactivated RSV-A2 and washed with PBS containing 0.05% Tween20 (PBST). Secondly, plasmas were diluted five times in PBST and incubated for 1 hour at room temperature while shaking. WHO RSV standard serum and adalimumab in 1% bovine serum albumin in PBST was used as a positive and negative control, respectively. Plates were then washed once with PBST, twice with PBS and twice with 200 µl ammonium bicarbonate. Elution of RSV-specific antibodies was performed by incubating with 100 mM formic acid for 5 minutes.

### Mass spectrometric IgG Fc-glycosylation analysis

Eluates containing either total or RSV-specific IgGs were collected in V-bottom plates, dried by vacuum centrifugation for 2.5 hours at 50°C and then dissolved in 20 µL RA buffer (0.4% sodium deoxycholate, 10 mM tris(2-carboxyethyl)phosphine (TCEP), 40 mM chloroacetamide, 100 mM tris(hydroxymethyl)aminomethane (TRIS) pH8.5) followed by shaking for 10 minutes and heating for 5 minutes at 95°C. Antibody samples were then subjected to proteolytic cleavage by adding 20 µL trypsin (10 ng/µL) and incubating for 20 hours at 37°C.

Analysis of IgG Fc-glycosylation was performed with nanoLC reverse phase (RP)-electrospray (ESI)-MS on an Ultimate 3000 RSLCnano system (Dionex/Thermo Scientific, Breda, The Netherlands) coupled to a Maxis HD quadrupole-time-of-flight MS equipped with a nanoBooster (Bruker Daltonics) using acetonitrile-doped nebulizing gas. (Glyco-)peptides were trapped with 100% solvent A (0.1% trifluoroacetic acid in water) and separated on a 4.5 min 3.0-21.7% solvent B (95% acetonitrile, 5% water) linear gradient (32). In the current study we focused on IgG1, without analyzing IgG3 due to its possible interference with IgG2 and IgG4 at the glycopeptide level (21). Mass spectrometry results were extracted and evaluated using Skyline software.

### Statistical analysis

Comparison of two groups or data points was performed by using a nonparametric Mann-Whitney test. Multiple comparisons were analyzed by using a nonparametric Kruskal-Wallis test, followed by Dunn’s multiple comparisons test. Multiple comparisons of categorical values were tested by Chi-square test. Correlations were assessed using a nonparametric Spearman’s rank-order correlation. Paired samples were analyzed using a nonparametric Wilcoxon matched-pairs test. *P* values <0.05 were considered statistically significant. Statistical analyses were performed and graphs were produced with Prism 8 software (GraphPad).

## Results

### Clinical characteristics of study subjects

Plasma samples were obtained from a total number of 84 infants below 7 months of age that were hospitalized for RSV infection (cases), hernia surgery (uninfected controls), or viral respiratory infections other than RSV (RSV-negative infected controls). Since we were primarily interested in the antibodies already present at the time of initial infection, we used RSV-specific IgA as a hallmark for newly elicited antibody production. Five children had IgA against at least 2 RSV antigens (AU/mL > 0.2) and were excluded (**Fig 1A**). In addition, we excluded 2 children that had presumably received Palivizumab, based on the fact that they displayed unusually high IgG levels for both pre-F and post-F, while G-specific levels were low or even below detection level.

**Figure 1.**
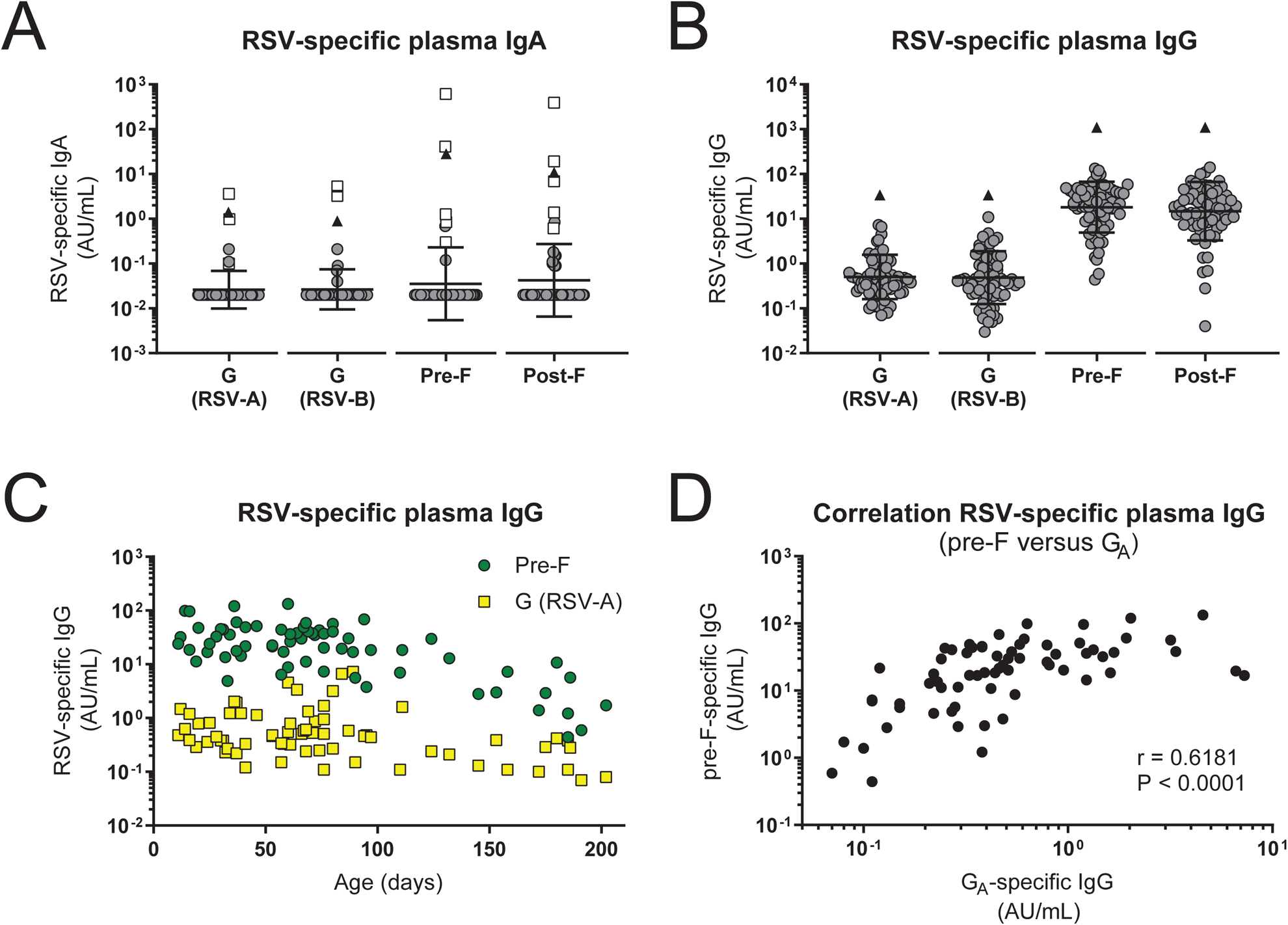
Characterization of the infant RSV-specific (maternal) antibody repertoire. An RSV-specific multiplex immunoassay was performed to quantify the concentration of IgA and IgG specific for RSV G (A- and B-strain), pre-F, and post-F in plasma. (A) Concentration of plasma IgA for each of the RSV antigens. Children that tested positive for IgA (AU/ml > 0.2) against a minimum of 2 RSV antigens are excluded for subsequent analyses (n=5, white squares). (B) Concentration of plasma IgG for each of the RSV antigens. Black triangles indicate the IgA/IgG levels of the adult reference serum pool. Since arbitrary concentrations were determined based on separate standard curves, concentrations cannot be directly compared between antigens. Graphs depict geometric mean and SD. (C) Concentration of RSV pre-F- and G_A_-specific plasma IgG plotted against age. (D) Correlation analysis of pre-F- and G_A_-specific maternal antibody levels. Nonparametric Spearman’s correlation analysis was used to assess correlations. Abbreviations: AU, arbitrary units; IgA, immunoglobulin A; IgG, immunoglobulin G; RSV, respiratory syncytial virus; SD, standard deviation.

The clinical and demographic characteristics of the children included in further analyses are depicted in **Table 1**. There was no significant difference in age, gestational age or breastfeeding between the three groups. However, there was a significant difference in sex between the three groups (P = 0.0459). A higher percentage of infants was infected with RSV-A (70%) compared to RSV-B (30%). The group of RSV patients contained both RSV mono-infections (n=28) and co-infections (n=15) with other respiratory viral pathogens, with rhinovirus being the most prevalent.

**Table 1.**
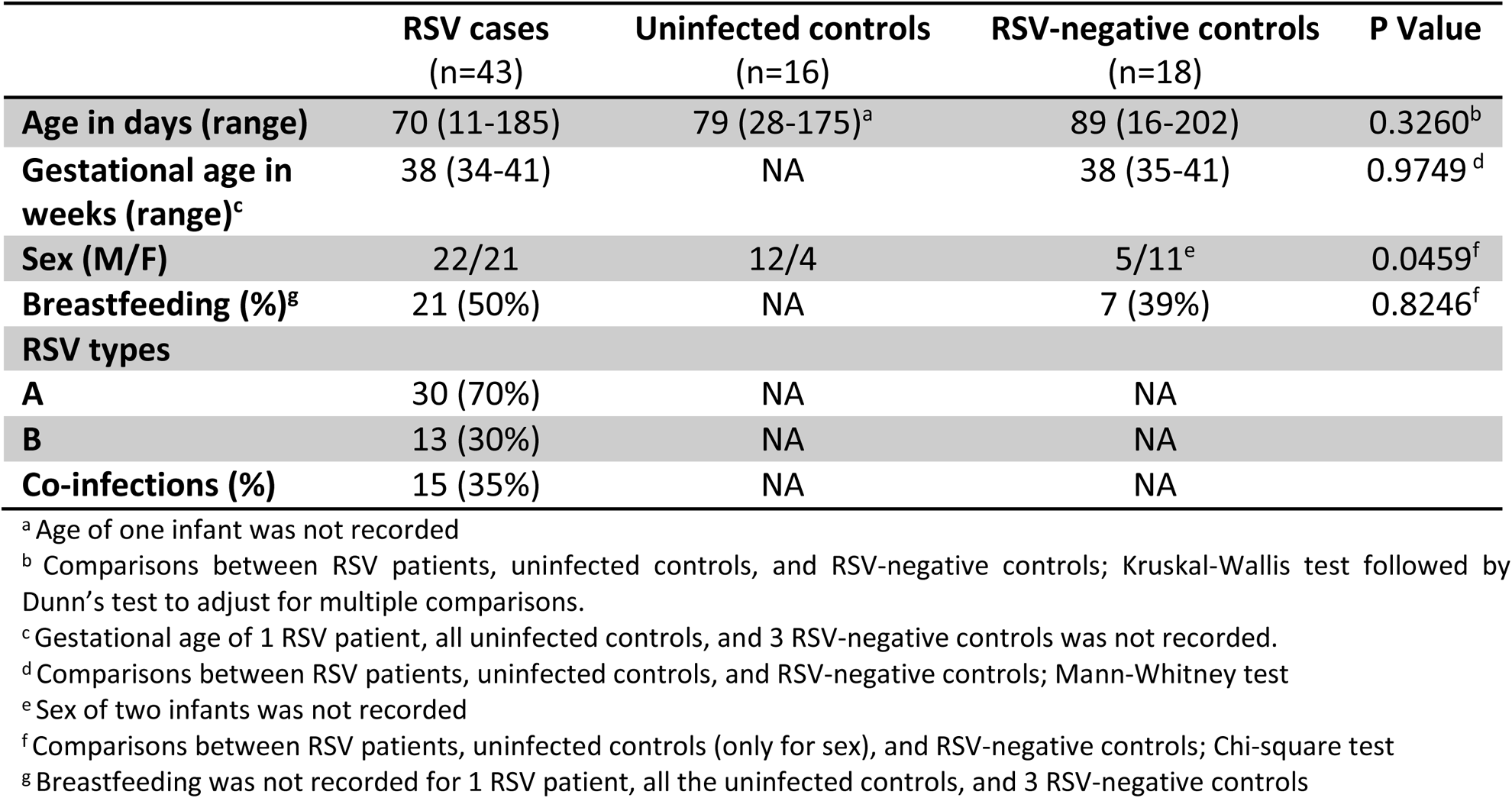
Demographic and clinical characteristics of study subjects.

### The concentration of RSV-specific IgG does not differ between RSV cases and controls

We used a multiplex immunoassay to quantify the concentration of IgG specific for the RSV attachment (G) and fusion (F) proteins. G protein from both an A- and B-strain was included, as these sequences are highly variable between the two RSV subtypes (33). Both the pre- and post-fusion conformations of the F protein were included, as these harbor partly overlapping but also distinct epitopes (34). We could detect antibodies to all four antigens in all samples and observed substantial spread between individuals (**Fig 1B**). Antibody levels were lower than those in the adult reference serum pool (triangles) in all samples. Despite the cross-sectional design of the study, we could observe a gradual waning of antibody concentrations with increasing age, as exemplified by antibodies targeting pre-F and G_A_ (**Fig 1C**). As expected, there was a positive correlation between the concentrations of antibodies targeting different RSV antigens, as exemplified for pre-F and G_A_ (**Fig 1D**). The concentrations of antigen-specific antibodies did not significantly differ between RSV-infected cases, uninfected controls, and RSV-negative infected controls (**Fig 2A-D**), confirming previous findings in a subset of this cohort (12).

**Figure 2.**
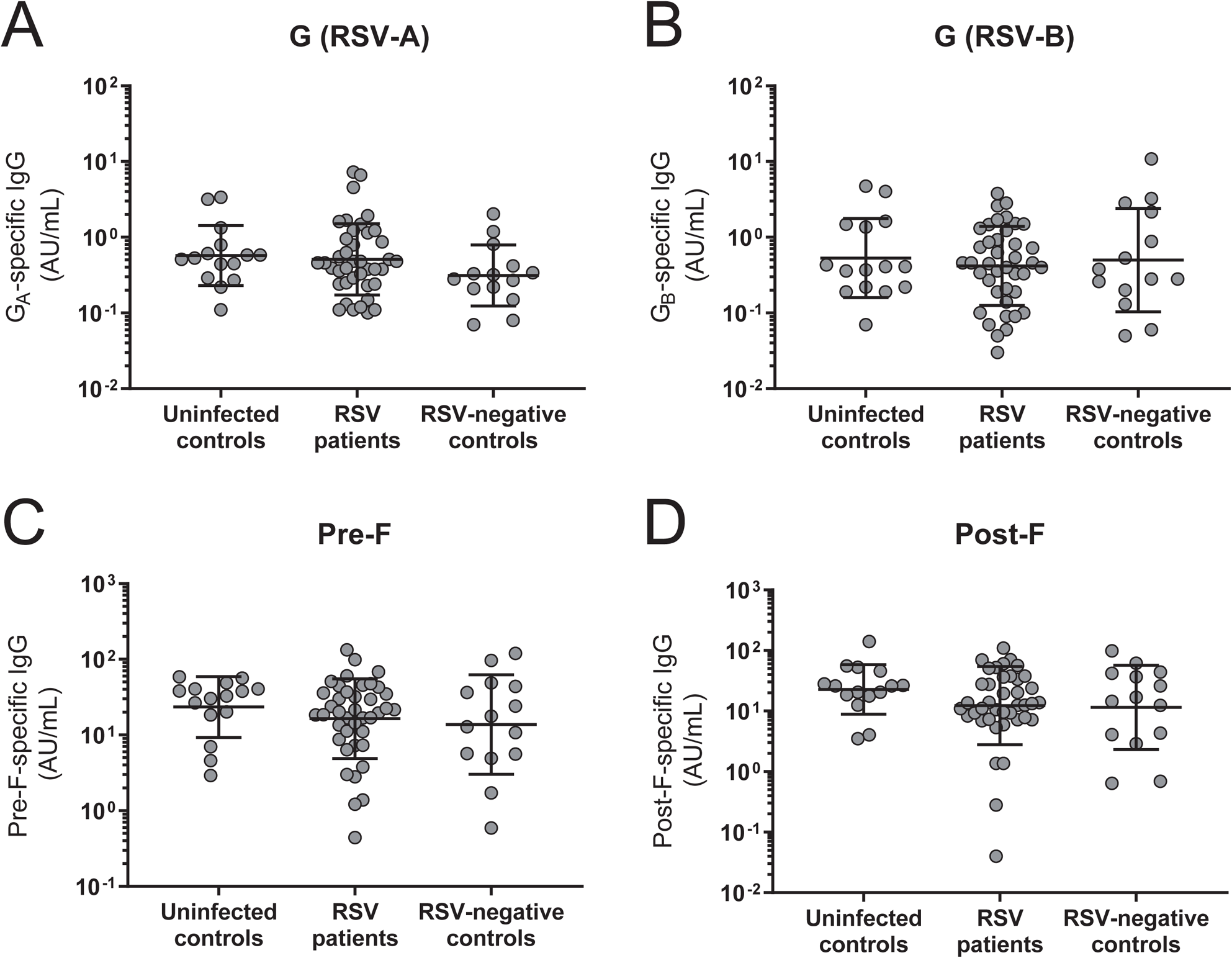
RSV-specific antibody concentrations do not differ between RSV cases and controls. An RSV-specific multiplex immunoassay was performed to quantify the amount of RSV antigen-specific IgG specific in plasma. Comparison of the antibody concentration against RSV-A attachment protein G (A), RSV-B attachment protein G (B), pre-fusion F protein (C), and post-fusion F protein (D) between uninfected controls, RSV patients, and RSV-negative infected controls. All graphs depict geometric mean and SD. Kruskal-Wallis test with Dunn’s multiple comparisons test was used for comparison between multiple groups, no significant differences were found. Abbreviations: AU, arbitrary units; IgG, immunoglobulin G; RSV, respiratory syncytial virus; SD, standard deviation.

### RSV-specific maternal antibodies activate NK cells *in vitro*

As NK cells play an important role in the control of viral lung infections, we next assessed whether RSV-specific maternal antibodies are capable of activating NK cells in an *in vitro* assay (schematically depicted in **Fig 3A**). To control for RSV-specificity, the assay was set up with a cord blood plasma pool that had been depleted for RSV-specific antibodies by incubation with RSV-infected cells, or with mock-infected cells as a control. Successful depletion was confirmed by multiplex immunoassay (**Fig 3B**). Incubation of NK cells with RSV-antibody complexes formed with a dilution series of mock-depleted cord blood plasma, naturally containing RSV-specific maternal antibodies, resulted in a concentration-dependent activation as measured by IFN-γ production (**Fig 3C**) and CD107a surface expression (**Fig 3D**), the latter being an established marker for NK cell degranulation (35). Importantly, cord blood plasma largely lacking RSV-specific antibodies (RSV-depleted) showed a strongly reduced NK cell activation.

**Figure 3.**
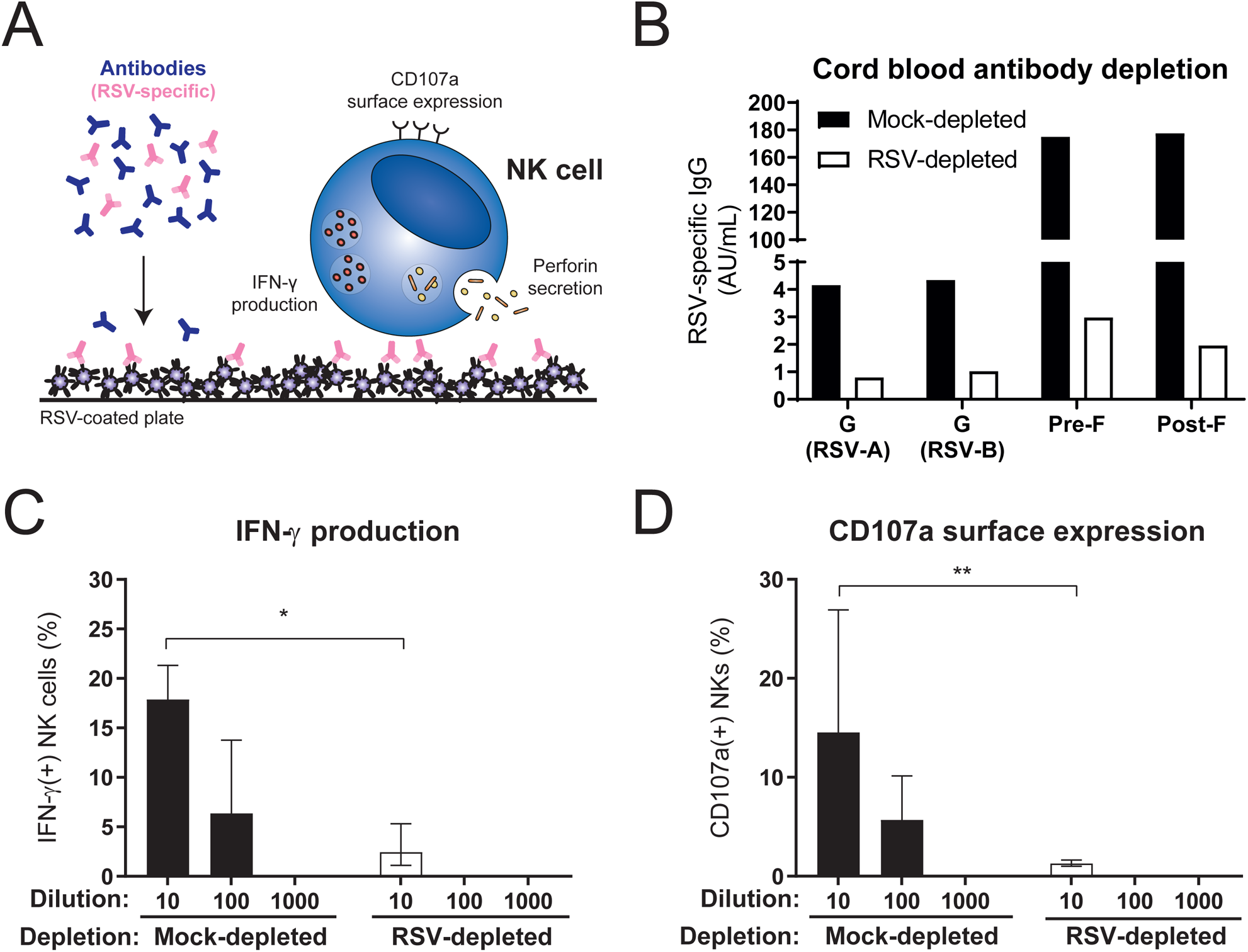
RSV-specific maternal antibodies activate NK cells *in vitro*. (A) Schematic representation of the NK cell activation assay. (B) RSV-specific multiplex immunoassay of mock-depleted (black) compared to RSV-depleted (white) cord blood plasma. NK cell-specific IFN-γ production (C) and CD107a surface expression (D) of three individual PBMC donors after 4 hour incubation with viral particles opsonized with mock-depleted or RSV-depleted cord blood plasma. Graphs depict geometric mean and SD. Kruskal-Wallis test with Dunn’s multiple comparisons test was used for comparison between mock- and RSV-depleted cord blood plasma with the same dilution (*P<0.05, **P<0.01). Abbreviations: AU, arbitrary units; CB, cord blood; IFN-γ, interferon gamma; NK cells, natural killer cells; RSV, respiratory syncytial virus.

### RSV-specific antibodies from infants with severe viral respiratory tract infections induce less IFN-γ production by NK cells than those from uninfected controls

Next, we assessed whether the capacity of RSV-specific antibodies to activate NK cells differed between RSV cases and controls. Since we observed considerable donor variation in primary NK cell activation in our initial experiments, we normalized the response for all individual PBMC donors (**Fig S2**). To this end, NK cell activation by the mock-depleted cord blood pool was set to 100 arbitrary units (AU). Activation by patient samples was expressed as AU relative to the mock-depleted cord blood pool for each individual PBMC donor. Finally, the overall response induced by each patient sample is obtained by averaging the normalized responses for all PBMC donors tested. Interestingly, we found that RSV-specific antibodies from both RSV patients and RSV-negative infected controls showed significantly decreased induction of NK cell IFN-γ production compared to uninfected controls (**Fig 4A**). Slightly less induction of CD107a expression was also observed for plasma from both RSV patients and RSV-negative infected controls compared to those from uninfected controls, but this difference was not statistically significant (**Fig 4B**). Since NK cell CD107a surface expression is used as a proxy for degranulation, we assessed its correlation with perforin secretion in the supernatant of the NK cell activation assay for one PBMC donor and found a strong positive correlation (**Fig 4C**). Finally, since the capacity of antibodies to activate NK cells is expected to be related to their concentration, we assessed the correlation between antibody levels and IFN-γ production. As exemplified for pre-F-specific antibody concentrations, there was only a moderate correlation with IFN-γ production (**Fig 4D**), suggesting that additional factors besides concentration contribute to the functional capacity of these antibodies.

**Figure 4.**
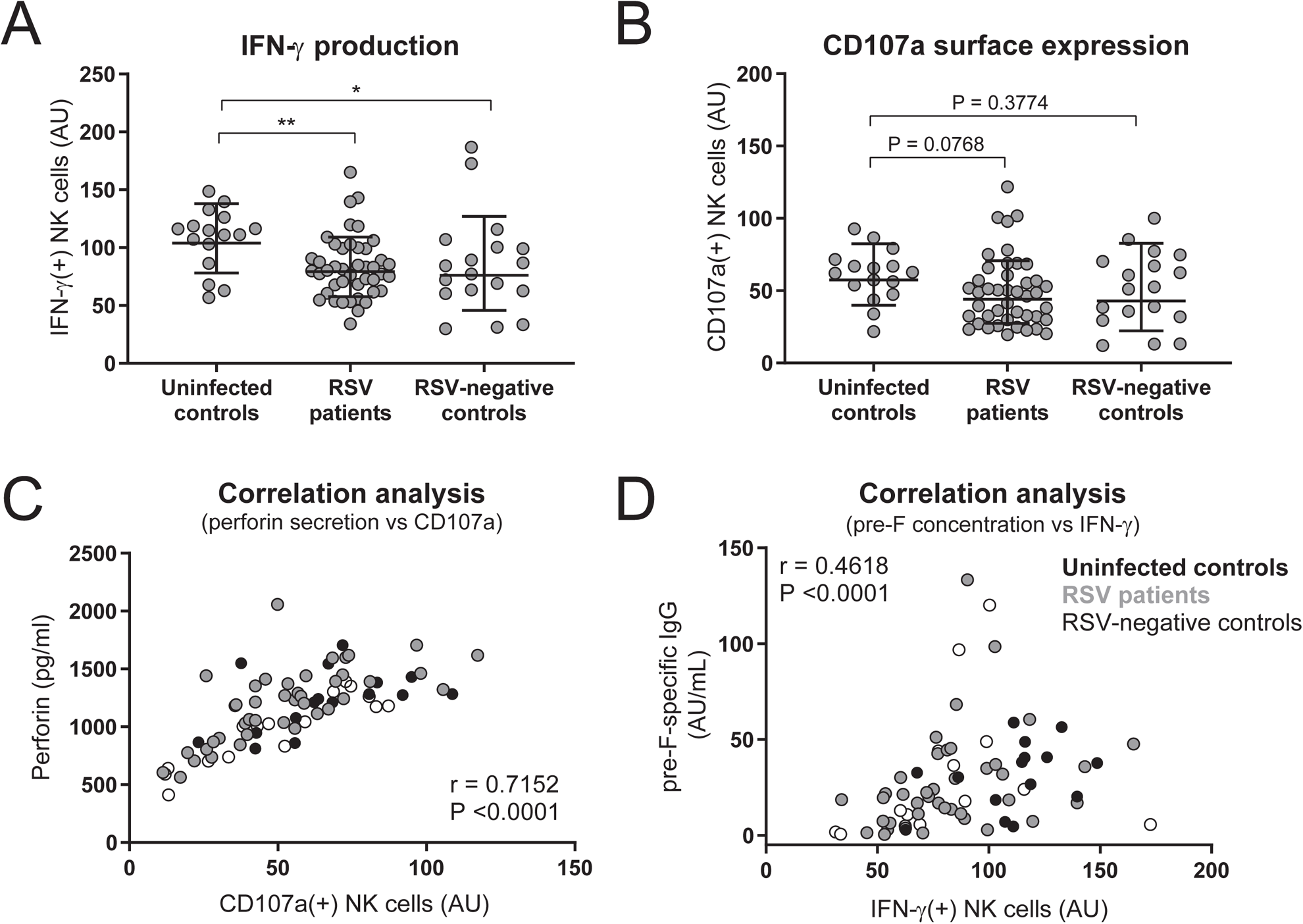
NK cell activation in RSV cases compared to controls. Normalized NK cell IFN-γ production (A) and CD107a surface expression (B) after 4 hour incubation of PBMCs (≤6 individual donors per sample) with viral particles opsonized with 10-fold diluted plasma samples from uninfected controls, RSV patients, or RSV-negative infected controls. Graphs depict geometric mean and SD. Kruskal-Wallis test with Dunn’s multiple comparisons test was used for comparison between multiple groups (*P<0.05, **P<0.01). (C) Correlation analysis between perforin secretion and CD107a surface expression of one donor. (D) Correlation analysis between RSV pre-F-specific antibody levels and IFN-γ production. Uninfected controls are indicated in black, RSV patients in grey, and RSV-negative infected controls in white. Nonparametric Spearman correlation analysis was used to assess correlations. Abbreviations: IFN-γ, interferon gamma; NK cells, natural killer cells; RSV, respiratory syncytial virus; SD, standard deviation.

### RSV-specific antibodies are differently glycosylated than total infant antibody pool

We next assessed the Fc-glycosylation status of total and RSV-specific antibodies in our cohort. To this end, we enriched samples for RSV-specific antibodies by adsorption to inactivated RSV-A2-coated plates and subsequent elution (schematically depicted in **Fig 5A**). Overall, we found a significantly higher abundance of galactosylation and sialylation in RSV-specific antibodies compared to total antibodies (**Fig 5B**). In addition, we noticed that for both RSV-specific and total antibodies the abundance of fucosylation increased with age (**Fig 5C-D**), while the abundance of galactosylation, sialylation, and bisecting *N-*acetylglucosamine (bisection) decreased with age (**Fig 5E-J**).

**Figure 5.**
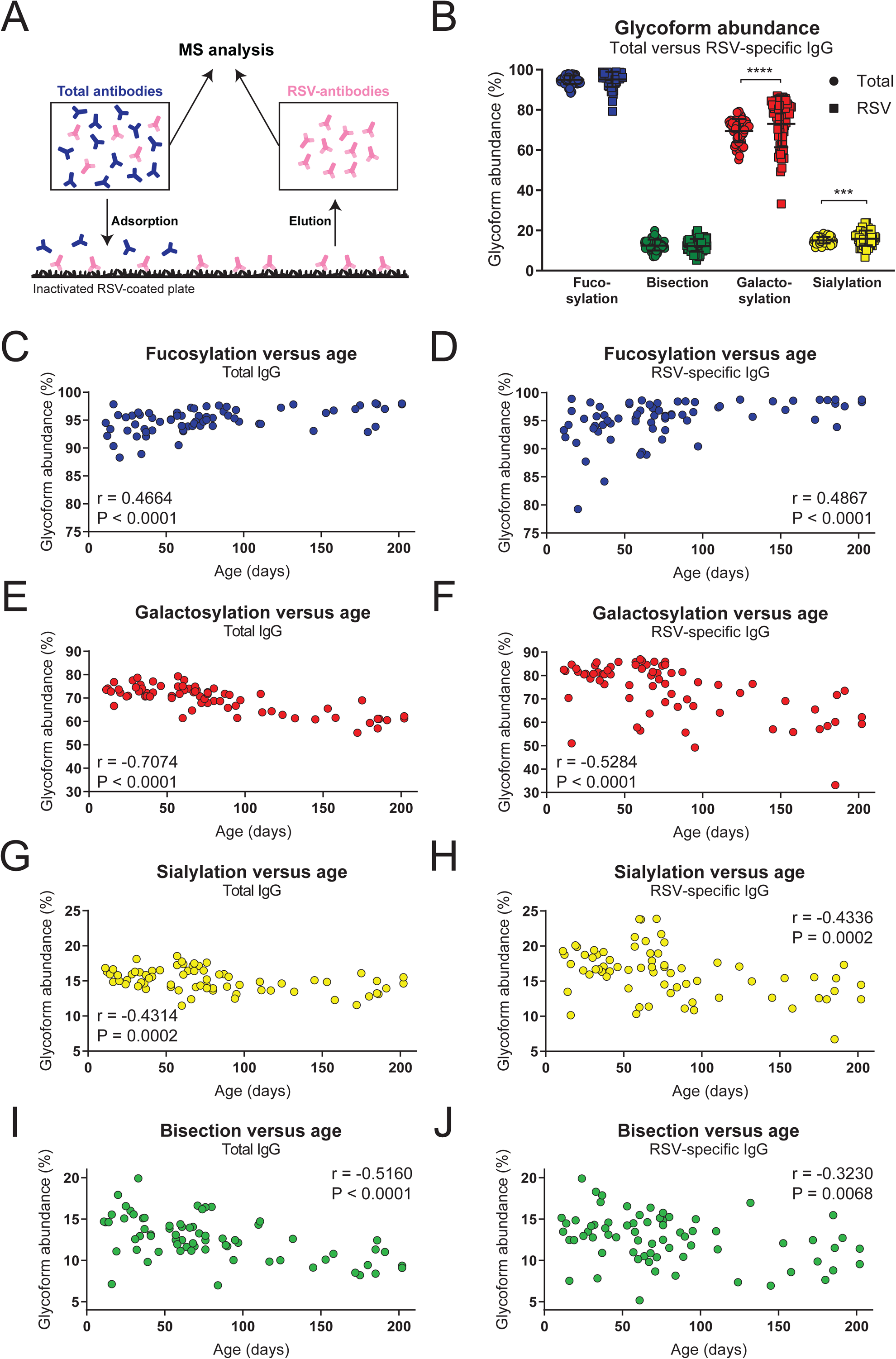
Fc-glycosylation pattern of total and RSV-specific antibodies. (A) Schematic representation of RSV-specific antibody enrichment. (B) Glycoform abundance of total compared to RSV-specific antibodies. Graph depicts geometric mean and SD. Nonparametric Wilcoxon matched-pairs test was used for comparison between total and RSV-specific antibodies (***P<0.001, ****P<0.0001). Correlation analysis between glycoform abundance (fucosylation, galactosylation, sialylation, bisection) and age for total antibodies (C, E, G, I) and RSV-specific antibodies (D, F, H, J). Nonparametric Spearman correlation analysis was used to assess correlations. AU, arbitrary units; IFN-γ, interferon gamma; NK cells, natural killer cells; RSV, respiratory syncytial virus; SD, standard deviation.

### Fucosylation of RSV-specific antibodies correlates with NK cell activation

The absence of fucose in the core Fc-glycan has been shown to increase binding affinity for FcγRIIIa, thereby increasing the capacity of the antibody for NK cell activation (23, 36). Similarly, galactosylation is known to enhance FcγRIIIa binding, albeit to a lesser extent than afucosylation, and this is particularly evident in the case of afucosylated IgG (23, 37). We therefore asked whether fucosylation of RSV-specific antibodies correlated with the antibody-induced NK cell activation observed in our assay. Indeed, although not very strong, we observed a significant negative correlation between fucosylation and the induction of NK cell IFN-γ production (**Fig 6A**) and CD107a surface expression (**Fig 6B**). Galactosylation of RSV-specific antibodies did not significantly correlate with IFN-γ production (**Fig 6C**), but it did show a small but significant positive correlation with CD107a surface expression (**Fig 6D**). Sialylation showed a small but significant positive correlation with both IFN-γ production (**Fig 6E**) and CD107a surface expression (**Fig 6F**). Bisection did not show a significant correlation with NK cell activation. As RSV-specific antibodies from infants with severe respiratory tract infections induced significantly less NK cell IFN-γ production compared to antibodies from uninfected controls, we asked whether this difference was reflected in the glycosylation status of these antibodies. However, none of the glycoforms showed a significant difference in abundance between cases and controls (**Fig 7A-D**).

**Figure 6.**
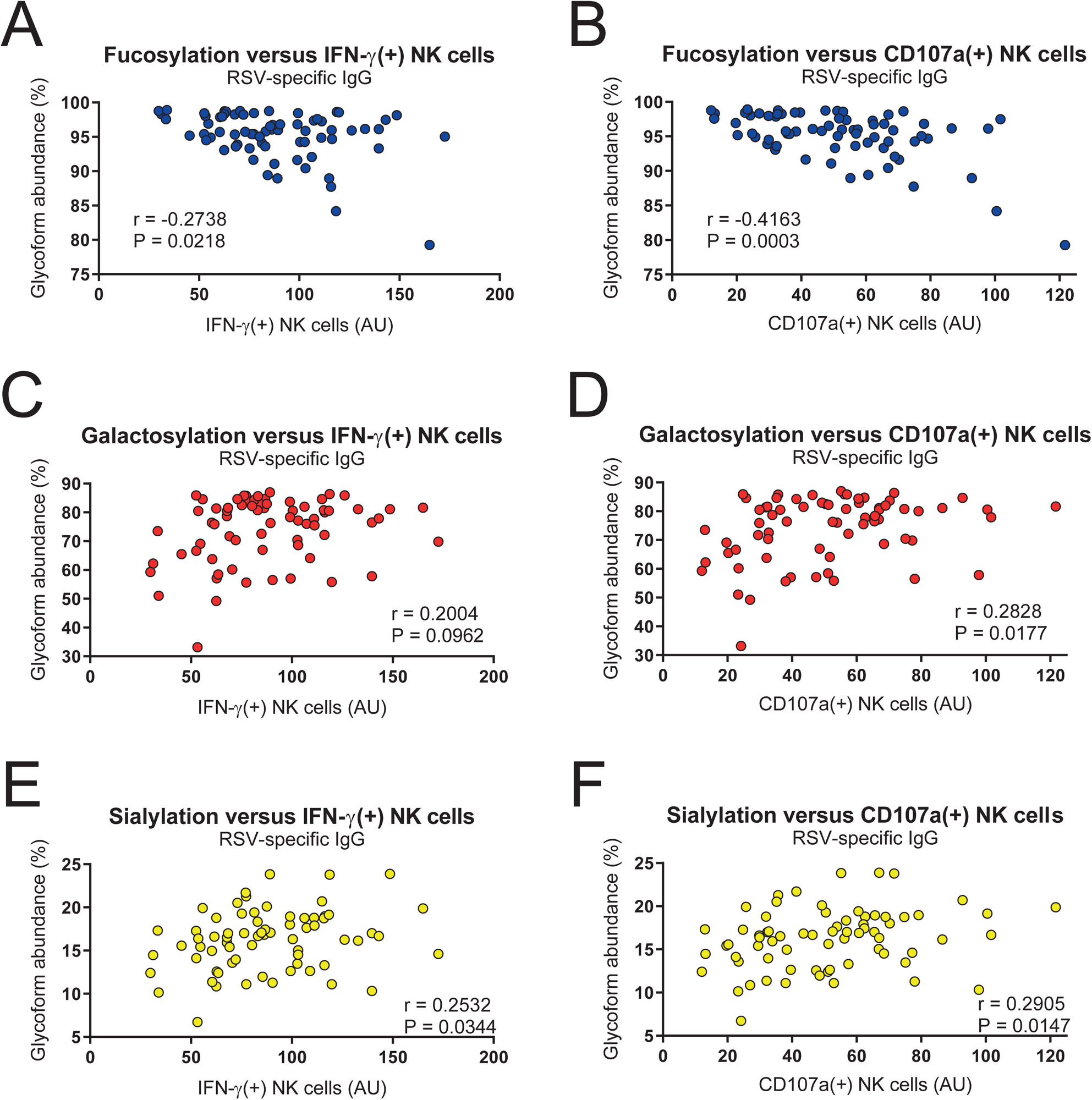
Fc-glycosylation correlates with NK cell activation. Correlation analysis between glycoform abundance of RSV-specific antibodies (fucosylation, galactosylation, and sialylation) and the capacity to induce IFN-γ production (A, C, E) and CD107a surface expression (B, D, F). Nonparametric Spearman correlation analysis was used to assess correlations. AU, arbitrary units; IFN-γ, interferon gamma; IgG, immunoglobulin G; RSV, respiratory syncytial virus.

**Figure 7.**
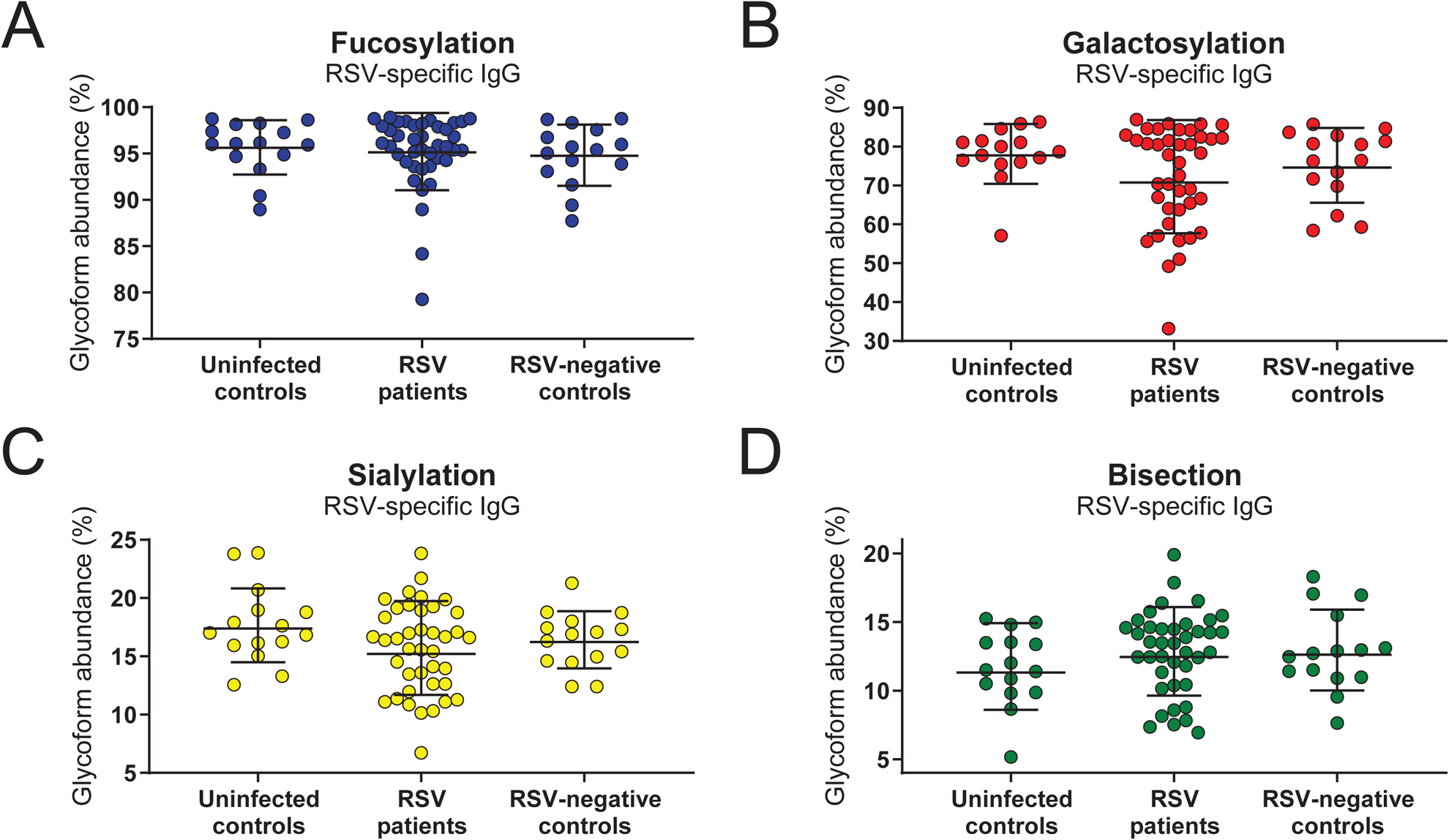
Fc-glycosylation does not differ between RSV cases and controls. Comparison of the abundance of fucosylation (C), galactosylation (D), sialylation (E), and bisection (F) between uninfected controls, RSV patients, and RSV-negative infected controls. Graphs depict geometric mean and SD. Kruskal-Wallis test with Dunn’s multiple comparisons test was used for comparison between multiple groups, no significant differences were found. IgG, immunoglobulin G; RSV, respiratory syncytial virus; SD, standard deviation.

## Discussion

Studies investigating the role of (maternal) antibodies in RSV infection mostly focus on neutralization and binding titers. Whereas antibody-mediated protection against RSV disease likely depends (at least in part) on virus neutralization, there is increasing evidence that Fc-mediated effector functions also play an important role during RSV infection (reviewed in (38, 39)). In two independent *in vivo* studies, modification of the Fc-domain of RSV-specific antibodies has shown major effects on lung viral titer (40, 41). These results strongly support the idea that Fc-dependent mechanisms contribute to the protective efficacy of RSV-specific antibodies. In the present study, we found that RSV-specific antibodies from infants hospitalized for RSV and/or other respiratory viral infections less potently induced NK cell IFN-γ production than those from uninfected controls. In addition, we found that NK cell activation significantly correlated with RSV antigen-specific antibody concentration and fucosylation, although neither of these factors significantly differed between cases and controls.

Antibody-mediated NK cell activation has gained increasing attention for its role in protection against a variety of infectious diseases. The capacity of antibodies to induce NK cell ADCC correlates with the control of mycobacterium tuberculosis (42) and the killing of chlamydia trachomatis (43). For HIV, ADCC-inducing antibodies have been identified as a key correlate of protection in the RV144 HIV vaccine trial (44-46). For influenza, the role of ADCC was shown to be protective in some studies (47, 48), but others point to the involvement of ADCC in exaggeration of the immune response (49-51). Taken together, these examples show that the capacity of pathogen-specific antibodies to engage NK cells can significantly contribute to the outcome of infection. Regarding RSV infection, in mice, NK cells have been shown to be sufficient to eliminate infection (52) and NK cell depletion was found to increase viral titers (53). However, NK cells have also been found to contribute to inflammatory lung injury in mice (53, 54). In humans, NK cells have been reported to be both decreased (55, 56) and increased (57, 58) compared to controls upon RSV infection. Thus, the exact role of NK cells during RSV infection remains to be established but our results suggest their activation contributes to a favorable outcome of infection.

We found that RSV-specific antibodies from both RSV-infected and RSV-negative infected controls showed a decreased propensity to activate NK cells compared to those from uninfected controls. Since we only have samples that were obtained during hospitalization, and not before infection, we cannot exclude that the difference we observe is due to the inflammation that these infants are experiencing. Although it has been shown that chronic infection and inflammation can affect functionality of the newly produced antibody repertoire (59), acute inflammation is unlikely to affect the antibodies that are already in circulation. A prospective study in which children are sampled before and during infection would provide more insight in the possible effects of inflammation.

Glycosylation analysis of the total antibody pool revealed changes related to infant age, where especially galactosylation showed a strong negative correlation with age. Fucosylation, on the other hand, showed a slight but significant increase with age. These observations are in line with the gradual replacement of maternal with infant antibodies. Altered glycosylation during pregnancy and preferential transplacental transfer result in increased IgG Fc-galactosylation of maternal antibodies (60-62). In addition, like all adult antibodies, maternal antibodies have lower fucosylation (94%) compared to highly fucosylated infant antibodies, which reach adult levels around the age of 20 years (60). We observed similar correlations between glycosylation and age for RSV-specific antibodies. This suggests that, despite the young age of the infants, our analysis is not entirely restricted to maternal antibodies. However, the increased galactosylation of RSV-specific antibodies compared to the total antibody pool, consisting of both infant and maternal antibodies, and the gradual waning of RSV-specific antibody concentrations seen with age suggest that we are looking primarily at maternal antibodies. However, this can only be firmly concluded when general glycosylation patterns of RSV-specific antibodies made by the infant are known.

Antibody functionality, and ultimately protective efficacy, is determined by a wide range of factors, including concentration, antigen and epitope specificity, glycosylation, isotype, and subclass. Each of these factors contributes to a greater or lesser extent to protection, which might explain why we did find a small but significant difference between cases and uninfected controls in antibody-mediated NK cell IFN-γ production, but not in CD107a surface expression, antibody concentration, and glycosylation. The multitude of factors contributing to antibody-mediated protection from severe RSV disease complicates the definition of a correlate of protection for this pathogen, which in turn hampers vaccine development, as efficacy has to be demonstrated in large-scale clinical trials. The use of a systems serology approach for RSV may offer an unbiased and comprehensive way to systematically investigate antibody characteristics and effector functions on a multidimensional level. It has already proven effective in identifying antibody features that contribute to protection against various other (viral) pathogens (42, 63, 64).

In conclusion, our results support the idea that antibody-mediated NK cell activation contributes to protection against severe RSV disease and warrant further investigations into the potential of harnessing Fc-mediated effector functions for improving the effectiveness of future RSV vaccines. A thorough understanding of the protective capacity, in its broadest sense, of (maternal) RSV-specific antibodies will be invaluable for the development of a safe and effective vaccine against this elusive pathogen.

## Acknowledgements

We want to thank Oliver Wicht for performing the cord blood depletion assay, Ronald Jacobi and Annelies Mesman for information regarding the NK cell activation assay protocol, and Lie Mulder for assistance with virus production. RSV post-F protein for the RSV multiplex immunoassay was kindly provided by Mark Esser from AstraZeneca.

## Figure legends

**Supplementary Figure 1. Gating strategy for NK cell activation assays.** Gating strategy for the measurement of CD107a surface expression and IFN-γ production in CD3(-)CD56(+) NK cells. Dot plots and histograms are depicted for one representative PBMC donor. Abbreviations: IFN-γ, interferon gamma; PBMCs, peripheral blood mononuclear cells; NK cells, natural killer cells.

**Supplementary Figure 2. Normalization of donor variation in NK cell activation assay.** NK cell-specific IFN-γ production (A) and CD107a surface expression (B) of six individual PBMC donors after 4 hour incubation with viral particles opsonized with the indicated concentrations WHO RSV reference serum. NK cell-specific IFN-γ production (C) and CD107a surface expression (D) after normalization to a mock-depleted cord blood plasma sample. All graphs depict geometric mean and SD. Abbreviations: AU, arbitrary units; IFN-γ, interferon gamma; NK cells, natural killer cells; RSV, respiratory syncytial virus; SD, standard deviation.

