## Supplementary figures and images for "Natural killer cell activation by respiratory syncytial virus-specific antibodies is decreased in infants with severe respiratory infections and correlates with Fc-glycosylation"

### Supplemental Figure 1

#### Supplemental Figure 1

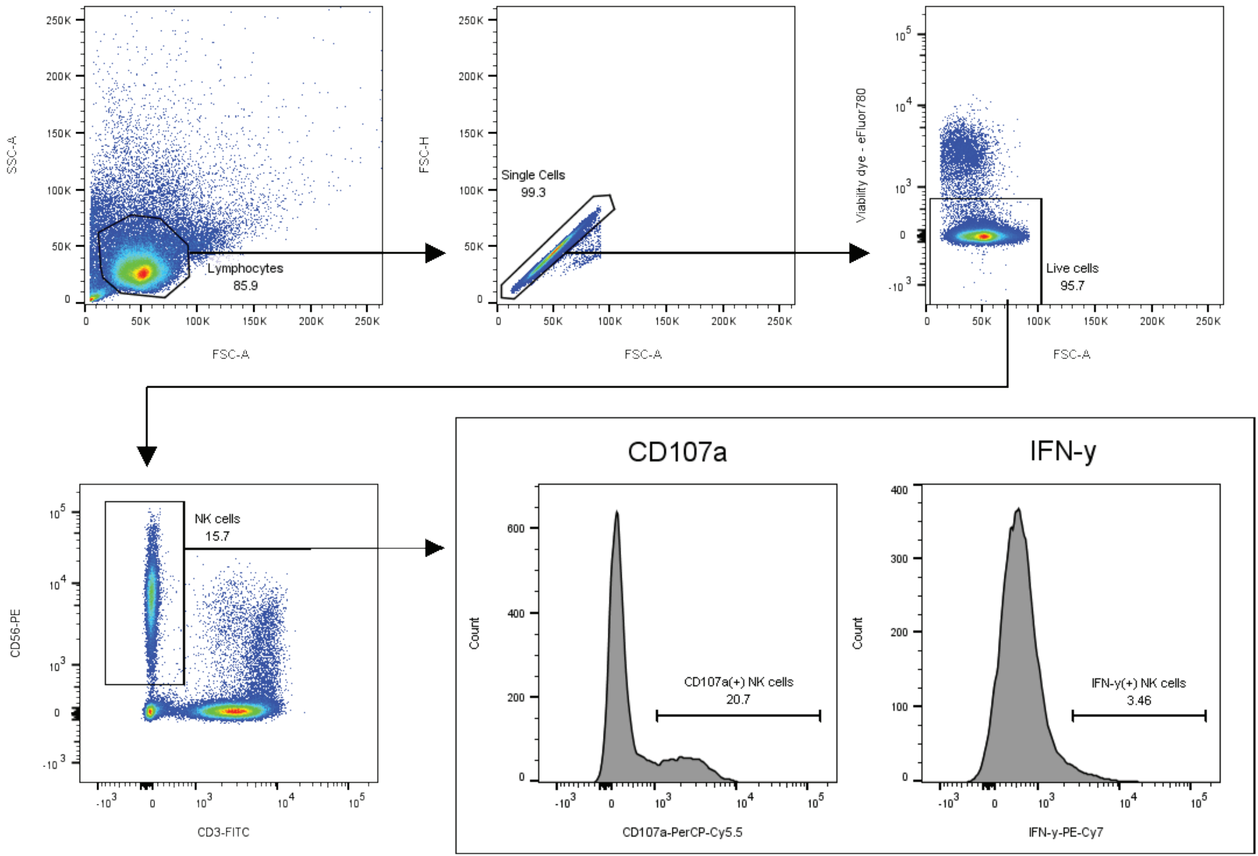

### Supplemental Figure 2

#### Supplemental Figure 2

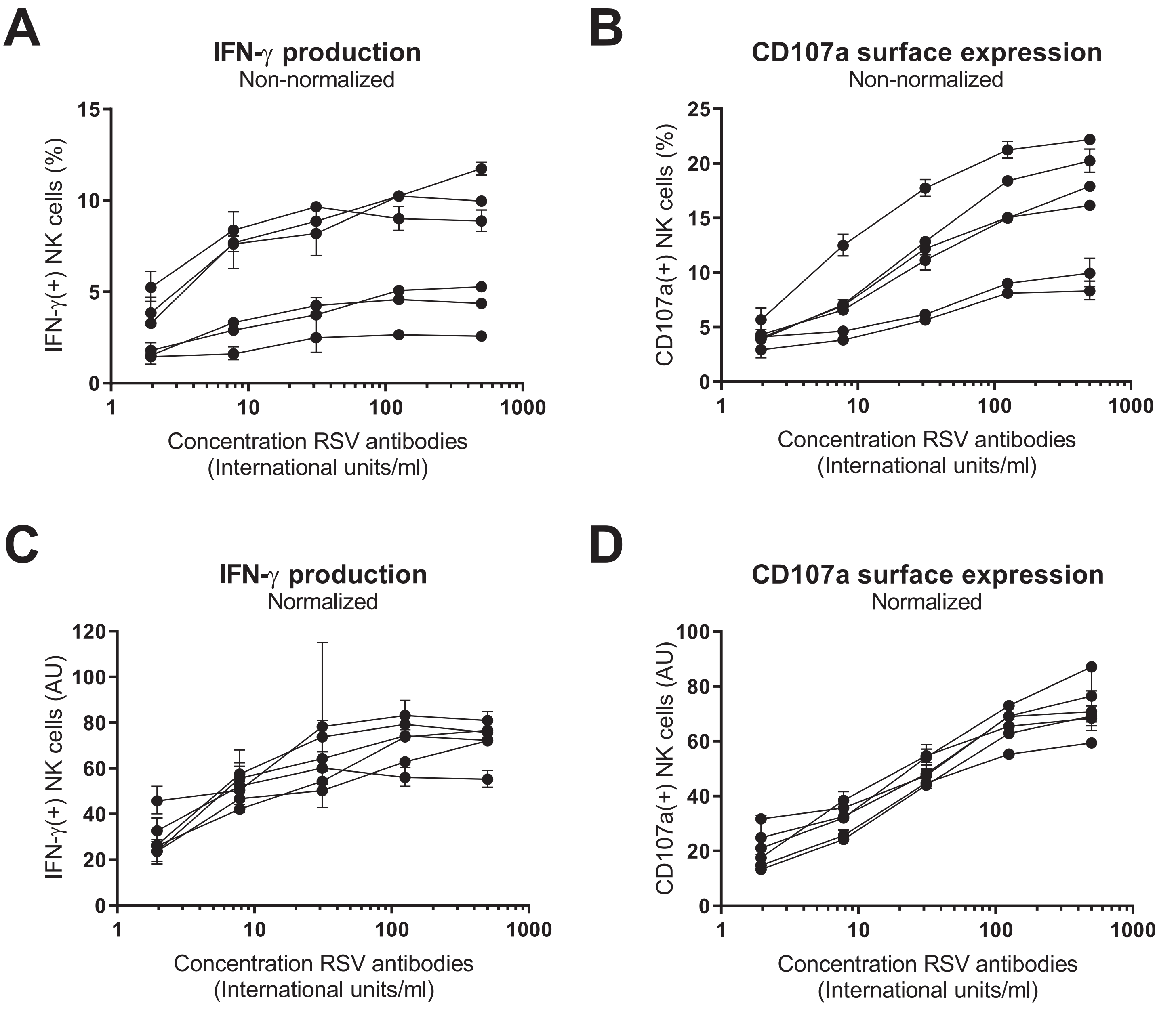
